# Cryo-EM Structure of *Salmonella typhimurium* ArnC; the Key Enzyme in Lipid-A Modification Conferring Polymyxin Resistance

**DOI:** 10.1101/2024.07.30.605912

**Authors:** Dhruvin H. Patel, Elina Karimullina, Yirui Guo, Cameron Semper, Deepak T. Patel, Tabitha Emde, Dominika Borek, Alexei Savchenko

**Author notes:** To whom correspondence should be addressed. Alexei Savchenko, Address: HSC B724 3330 Hospital Drive NW, Calgary, AB. Equal contribution (Elina Karimullina, Yirui Guo).

## Abstract

Polymyxins are last-resort antimicrobial peptides administered clinically against multi-drug resistant bacteria, including Gram-negative ESKAPE pathogens. However, an increasing number of pathogens employ a defense strategy involving a relay of enzymes encoded by the *pmrE(ugd)* loci and the *arnBCDTEF* operon. As a result, an Ara-4N headgroup is added to the lipid-A component of outer membrane (OM) lipopolysaccharides (LPS) rendering polymyxins ineffective. Here, we report the cryo-EM structures of glycosyltransferase ArnC from *Salmonella typhimurium* resolved in both apo and UDP-bound forms at resolutions 2.75 Å and 3.8 Å, respectively. The structure of the ArnC protomer comprises of three distinct regions: an N-terminal glycosyltransferase domain, transmembrane region, and the interface helices (IHs). ArnC forms a stable tetramer with C2 symmetry through interactions in the C-terminal region, which is expected to protrude into the cytosol, where the β8 strand inserts into the adjacent protomer. ArnC protomers have two distinct types of interfaces involving multiple hydrogen bonds and salt bridges. The binding of UDP induces conformational changes that stabilizes structurally labile A-loop, spanning residues 201 to 213, and part of the putative catalytic pocket formed by IH1 and IH2. The comparative analysis of ArnC structures with homologs GtrB and DPMS suggests the key residues involved in ArnC catalytic activity.

## 2. Introduction

Polymyxins are used as last-resort antibiotics against multi-drug resistant bacterial pathogens, including *Pseudomonas aeruginosa* and *Salmonella typhimurium*. These naturally derived lipopeptides, which include clinically administered polymyxin B and colistin, share a conserved scaffold, consisting of a cationic polypeptide ring and a hydrophobic acyl chain (1–3). When in contact with Gram-negative bacteria, the polypeptide ring electrostatically binds to the phosphate groups of lipid A moiety of lipopolysaccharides (LPS), which forms the outer leaflet of the outer membrane (OM) in these organisms. Subsequently, the polymyxin acyl chain perforates the outer membrane, leading to leakage of periplasmic proteins and further intake of polymyxins, which ultimately leads to bacterial cell death (4).

Bacteria have developed resistance to polymyxins through various mechanisms, including LPS modifications that result in membrane rigidification and charge alteration (5). Among the most prevalent LPS modifications is the addition of 4-amino-4-deoxy-L-arabinose (L-Ara4N) sugar to the lipid-A phosphate headgroup, which reduces the net negative charge of the outer membrane, and thus, prevents polymyxin binding. Inhibiting this lipid-A modification restores bacterial susceptibility to cationic antimicrobial peptides, including immune peptides and polymyxins (5).

The amino arabinose biosynthetic pathway, which mediates the addition of L-Ara4N, consists of enzymes encoded by the *pmrE(ugd)* and the *arnBCDTEF* operon (6–8). Starting with the glucose-UDP (uridine diphosphate) as the substrate, the enzymatic activities of Ugd, ArnA, and ArnB result in the synthesis of UDP-4-deoxy-4-formamido-beta-L-arabinopyranose (UDP-L-Ara4FN) in the bacterial cytosol (9–13). UDP-L-Ara4FN is then transferred by the integral membrane protein ArnC (undecaprenyl-phosphate 4-deoxy-4-formamido-L-arabinose transferase) to undecaprenyl phosphate (UndP), generating UndP-L-Ara4FN in the inner membrane (10). Next, ArnD deformylase catalyzes hydrolysis of UndP-L-Ara4FN to UndP-L-Ara4N. UndP-L-Ara4N is then transported to the outer membrane where ArnT catalyzes the attachment of the L-Ara4N group to the 4’-phosphate of core lipid-A (10, 14). Given their importance in developing polymyxin resistance, proteins from this pathway have been characterized extensively, with the exception of ArnC.

ArnC is localized to the inner membrane and is classified as a type-2 glycosyltransferase (GT-2) based on sequence similarity (15). In polymyxin-resistant *E. coli* carrying the *arn* operon, the deletion of *arnC* gene decreased the level of UndP-Ara4FN, and the deletion of the adjacent *arnD* gene led to the accumulation of UndP-Ara4FN, confirming the role of ArnC in the formation of UndP-Ara4FN (16). However, how ArnC performs its function has not been characterized at the molecular level. Therefore, we used cryogenic electron-microscopy single particle reconstruction (cryo-EM SPR) to determine the structures of *Salmonella typhimurium* ArnC in apo and UDP bound forms.

## 3. Materials and Methods

### 3.1 Protein expression and purification

A synthetic gene (TWIST DNA) encoding *Salmonella spp.* ArnC was codon optimized and cloned into pMCSG53 for overexpression in *E. coli* (6x-His-ArnC). A previously described protein expression and purification protocol was followed, but with slight modifications (17).

The vector was transformed into *E. coli* C43 (DE3) strain, which was then grown in Luria broth (LB) supplemented with ampicillin (100 μg/mL) to an OD600 of 0.55 at 37 °C. The cultures were then incubated on ice for 1 hour, induced with 1 mM isopropyl-β-d-thiogalactopyranoside (IPTG), and grown overnight at 20 °C. All further steps were conducted at 4 °C unless specified otherwise.

Cells were collected by centrifugation at 5000×g for 20 minutes and were resuspended in 1×PBS buffer pH 7.4. The cell suspension was supplemented with phenylmethylsulfonyl fluoride (PMSF, 0.5 mM), DNase1 (20 μg/mL), lysozyme (1 mg/mL), and TCEP (0.5 mM), followed by three freeze-thaw cycles. The lysed cells were homogenized using a cell disruptor (Avastin) at 15,000 psi and the cell debris were separated by centrifugation (13,500×g for 20 minutes at 4°C). The supernatant was centrifuged at 40,000 rpm for 1 hour 40 minutes to extract the crude membrane fraction, which was then washed and ultracentrifuged in 1× PBS solution at 40,000 rpm for 1 hour and 30 minutes, and the isolated membrane was stored at −80 °C.

To solubilize 6x-His-ArnC, the isolated membrane was resuspended in extraction buffer overnight (50 mM Na Phosphate pH 7.6, 300 mM NaCl, 20% [v/v] glycerol, 0.5 mM TCEP), and then incubated with 1% n-dodecyl-β-D-maltoside (DDM) for 2 hours. The membrane extract suspension was ultracentrifuged at 40,000 rpm for 40 minutes, and the supernatant was incubated with Ni-NTA beads in 50 mM imidazole for 18 hours. ArnC bound Ni-NTA beads were collected and washed in a gravity column with wash buffer 1 (50 mM Na phosphate pH 7.6, 500 mM NaCl, 15% [v/v] glycerol, 0.5 mM TCEP, 40 mM imidazole, 0.1% [w/v] DDM) and then wash buffer 2 (50 mM Na Phosphate pH 7.6, 300 mM NaCl, 10% [v/v] glycerol, 0.5 mM TCEP, 50 mM imidazole, 0.05% [w/v] DDM). 6x-His-ArnC was eluted with elution buffer (50 mM Na phosphate pH 7.6, 300 mM NaCl, 5% [v/v] glycerol, 0.5 mM TCEP, 0.05% [w/v] DDM, 300 mM imidazole), and dialyzed overnight with TEV protease in dialysis buffer (50 mM Tris pH 8.0, 300 mM NaCl, 5% [v/v] glycerol, 0.5 mM TCEP, 0.05% [w/v] DDM). After passing the dialyzed eluent through the Ni-NTA resin, the His-tag free ArnC flow-through fractions were collected and stored at −80 °C. To enhance protein stability, ArnC was incubated with amphipol A8-35 (Anatrace) in a 1:3 protein-to-amphipol ratio for 4 hours, followed by incubation with Bio-beads (Biorad) for 16 hours. Bio-beads were removed by passing the solution through an empty column. Amphipol stabilized ArnC was concentrated using an Amicon concentrator and then applied to a Superdex 200 Increase 10/300 column (Cytiva) equilibrated in cryogenic buffer (50 mM Tris pH 8.0, 150 mM NaCl, 0.5 mM TCEP). Fractions containing ArnC of the highest purity were concentrated for structural studies.

### 3.2 Grid preparation and data collection

Quantifoil R 1.2/1.3 300 Mesh Gold grids were glow-discharged for 90 s at 30 mA using a PELCO easiGlow™ system to obtain a hydrophilic surface. Subsequently, 3 μL of a particular ArnC sample was applied to the grid surface under 100% humidity at 4°C, followed by blotting for 5.0-5.5 s with a blot force of 18 or 19, and cryo-cooling in liquid ethane using a Thermo Scientific VitrobGt Mark IV. The image data were acquired with the beam-image shift method on a Titan Krios G2 microscope equipped with a K3 Summit direct electron camera operated in super-resolution mode with the GIF Quantum Energy Filter’s slit width set to 25 eV. Both data sets were collected at a nominal magnification of 105,000× (physical pixel size of 0.834 Å), with a defocus range of −1.0 to −3.0 μm, and 3×3 movies acquired per stage movement with SerialEM. A phase plate and objective aperture were not used. Movies were dose-fractionated into 125 frames with a total dose of 100 e^-^/Å^2^. For the UDP+ArnC sample, we mixed 20 µl of the ArnC sample with UDP and MnCl2, each at a 10 mM stock solution, to a final concentration of 0.5 mM for both. The mixture was incubated for ∼15 minutes on ice before grid preparation. The 10 mM stock solution of UDP was taken from the UDP-Glo™ Glycosyltransferase Assay (Promega, Part #V698A).

### 3.3 Cryo-EM image processing and model building

8,583 movies were collected for ArnC apo. 4,353 movies were collected for ArnC+UDP+Mn^2+^. All movies were imported into CryoSPARC followed by patch motion correction using binning of 2 and patch CTF correction (18). Particles were first picked using blob picker on all micrographs to be used as training datasets for TOPAZ particle picker (19). Top micrographs (215 for ArnC apo, 86 for ArnC+UDP+Mn^2+^) with the highest number of particles picked were used as input for TOPAZ training to further enrich the particles. Picked particles were extracted with a box size of 360 pixels and cleaned by several rounds of 2D classifications (450,749 particles for ArnC apo, 95,489 particles for ArnC+UDP+Mn^2+^).

*Ab initio* reconstruction was performed on the clean particle stack (4 classes for ArnC apo, 3 classes for ArnC+UDP+Mn^2+^) to separate ArnC tetramer from broken particles and contaminants. The classes with pure ArnC tetramer (311,189 particles for ArnC, 58,569 particles for ArnC+UDP+Mn^2+^) were selected for homogeneous refinement and non-uniform refinement in C2 symmetry and resulted in maps with a resolution of 2.75 Å and 3.8 Å for ArnC apo and ArnC+UDP+Mn^2+^, respectively (18).

To visualize the motion of the ArnC apo, a C2 symmetry expansion was applied to the clean particles used to reconstruct the 3D map. A mask that excludes the amphipol ring around the transmembrane domain was also generated. This mask and the symmetry-expanded particles were used as inputs for the 3D Variability job in CryoSPARC (18). Three modes were solved in the analysis with C1 symmetry and filter resolution of 6 Å. Results were exported using 3D Variability Display job (simple mode) with 20 frames per mode. Movies for each mode/component were generated by Chimera (20).

An initial atomic model was obtained by manually docking the Alphafold2 model of the ArnC tetramer to the map using Coot (21). The model was then iteratively rebuilt manually in Coot and refined using Phenix.real_space_refine and Servalcat (21–27). Model statistics can be found in Table 1 and workflow is summarized in Figure S1. The atomic coordinates (PDB 9BHC for ArnC apo, PDB 9BHE for ArnC+UDP+Mn^2+^) and maps (EMD-44540 for ArnC apo and EMD-44542 for ArnC+UDP+Mn^2+^) have been deposited in the Protein Data Bank (http://wwpdb.org/) and the Electron Microscopy Data Bank (https://www.ebi.ac.uk/emdb/), respectively.

**Table 1.**
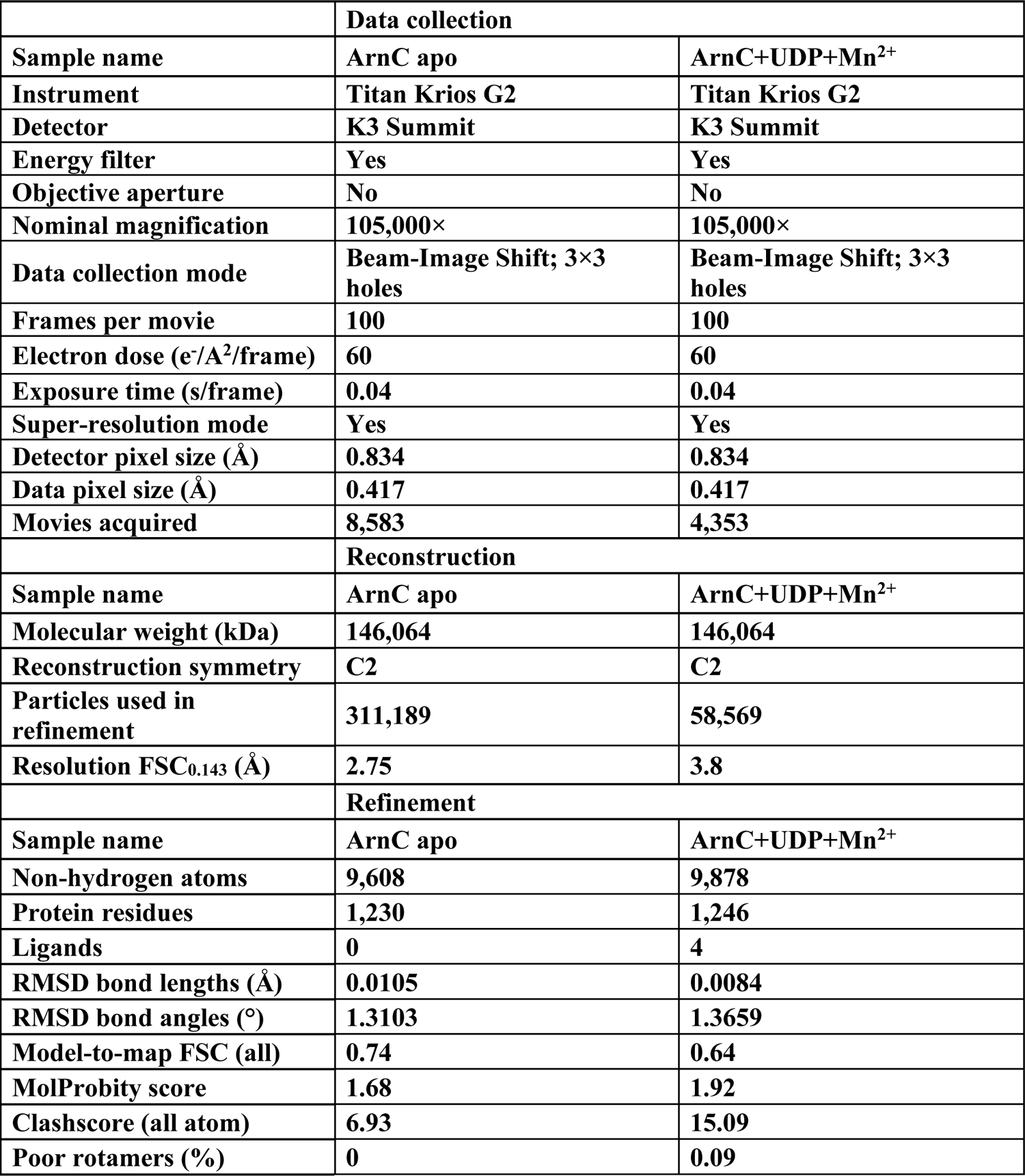
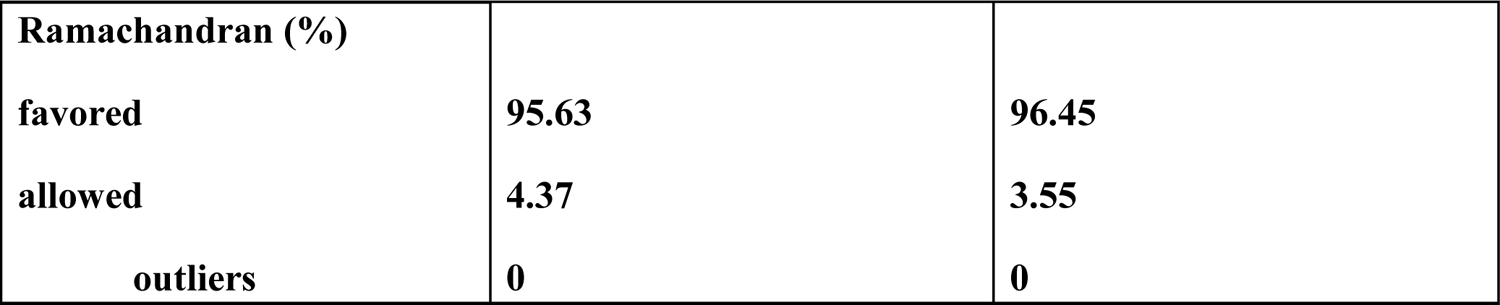
Data collection and processing.

## 4. Results and Discussion

### 4.1 Structure of ArnC Protomer

The overall structure of the ArnC protomer follows the general architecture observed in other membrane-bound glycosyltransferases and consists of three regions (Figure 1A). The N-terminal region (residues 1 to153) folds into a Rossman-like α-β domain that is similar to the canonical GT-A domain common for the GT-2 family of glycosyltransferases. This domain in the ArnC structure is followed by two α-helices (α6 and α9) positioned along the expected plane of the membrane. These helices have been colloquially referred to as juxtamembrane/interface helices and are henceforth referred to as interface helix 1 (IH1) (residues 134 to 153) and interface helix 2 (IH2) (residues 213 to 229) (Figure 1B). These helices are near parallel to one another with a slight tilt and are amphipathic in nature with the surface facing the membrane enriched with hydrophobic residue. The C-terminal portion of ArnC contains two transmembrane (TM) helices, which we will refer to as TM1 (residues 232 to 256) and TM2 (residues 267 to 304). TM1 is immediately downstream of IH2 and positioned perpendicular to the IH1/IH2 plane. A short periplasmic loop formed by residues 262 to 270 connects TM1 to TM2, with the end of TM2 expected to protrude outside the membrane and into the cytoplasm. The TM2 is followed by a C-terminal linker, termed as C-linker, (residues 305 to 309) that leads to the C-terminal β8 strand (residues 310 to 313) in the cytoplasm that points away from the originating protomer.

**Figure 1:**
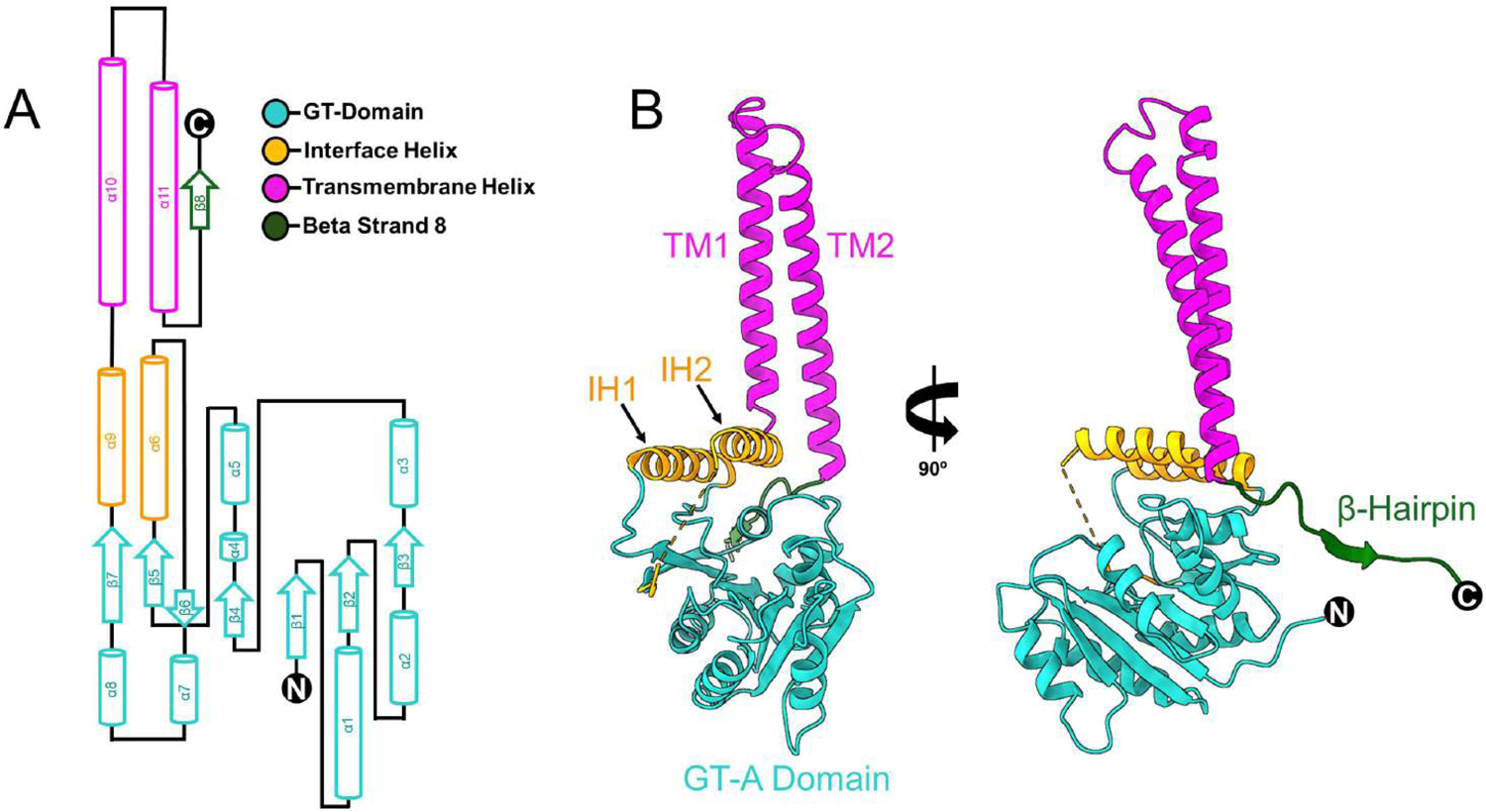
Structure of ArnC protomer (A) Topological arrangement and secondary structure elements of the ArnC protomer (B) Ribbon representation of the ArnC protomer.

### 4.2 Structure of Apo ArnC tetramer

ArnC forms a stable tetramer with C2 symmetry, which buries 19,000 Å² out of 51,000 Å² of the total surface area. In this quaternary structure, the C-terminal β8 strand of each protomer which follows the TM2 inserts into the N-terminal domain of an adjacent protomer, integrating with its structure. The outer surface of the cytosolic domains is hydrophilic and has anionic charge, but a positively charged pocket is observed at the IH1/2 surface facing the GT-2 domain (Figure 2B).

**Figure 2:**
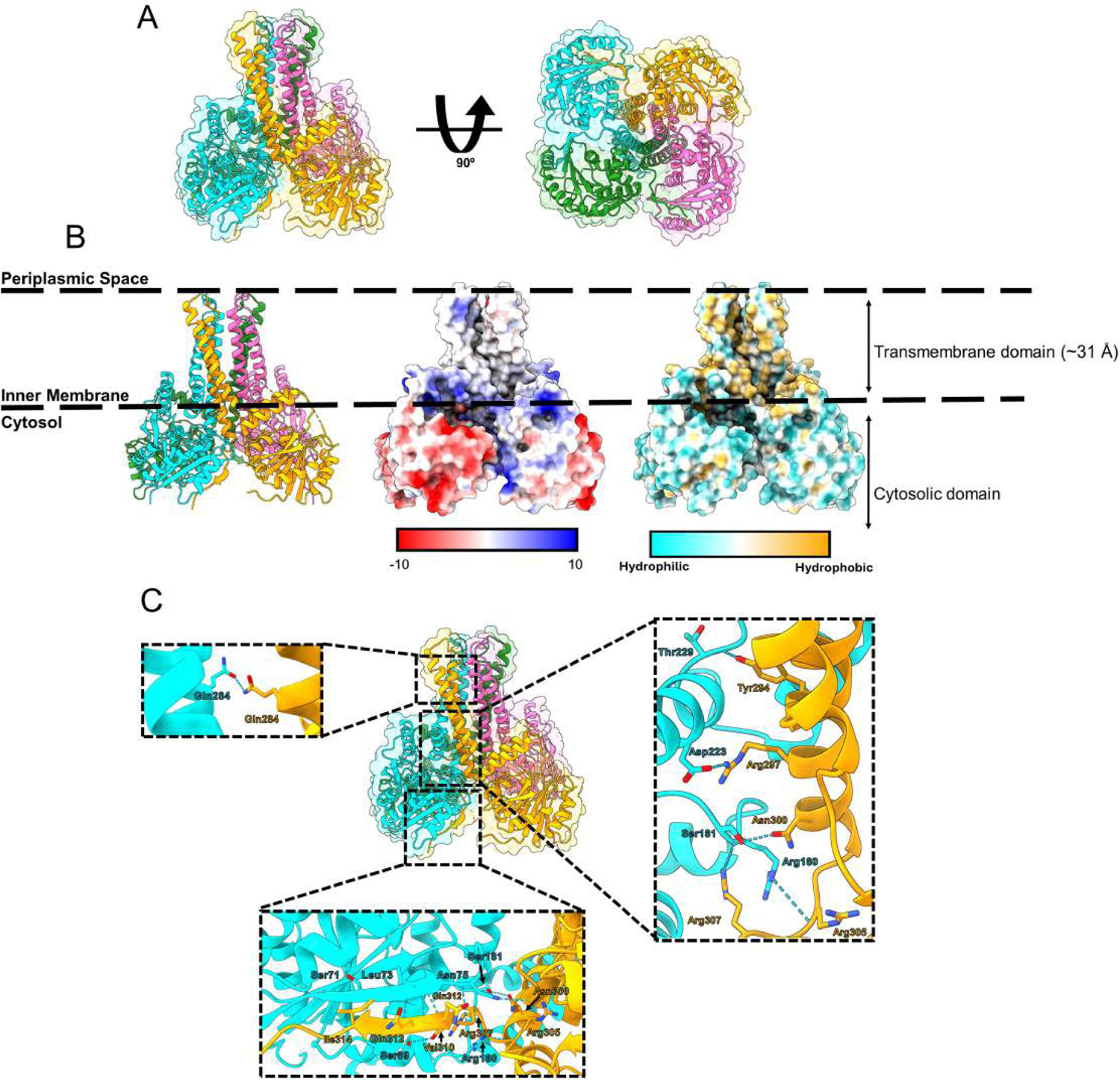
Structure of ArnC tetramer (A) Cytoplasmic view of the ArnC Tetramer. Protomer are color coded; A: tangerine, B: pink C: green D: teal (B) Biophysical properties of the ArnC tetramer modelled in the membrane. The depth of the TM in the membrane was computed using the PPM server (31) (C) Residues involved in inter-subunit hydrogen bonding in the tetrameric quaternary structure.

In addition to β8 strand swap, the ArnC tetramer is stabilized by extensive interactions between adjacent protomers, while minimal contacts were observed between diagonally located protomers (Figure 2A). There are two types of interfaces formed by each protomer in the ArnC tetramer. Interface type I is formed between protomer D and protomer A in D-A dimer, and protomer C and protomer B in B-C dimer. This interface buries an average of 2366 Å² and involves interactions between120 residues, which includes 26 hydrogen bonds and 2 salt bridges. Interface type II, observed between protomer B and D (B-D dimer), and protomer A and C (A-C dimer), buries an average of 2345 Å². Similar to interface type I, approximately 120 residues are involved in intermolecular interactions, which include 21 hydrogen bonds and 3 salt bridges (Table 2).

**Table 2.** Summary of ArnC tetramer interface statistics (33, 34).

| **Type** | **Chains** | **No. of**<br>**interface**<br>**residues** | **Interface Area**<br>**(Å<sup>2</sup>)** | **No. of hydrogen**<br>**bonds** | **No. of salt**<br>**bridges** |
| --- | --- | --- | --- | --- | --- |
| **1** | **D-A** | **70:52** | **2377** | **26** | **2** |
| **1** | **B-C** | **69:52** | **2355** | **25** | **2** |
| **2** | **B-A** | **52:66** | **2347** | **21** | **3** |
| **2** | **D-C** | **51:66** | **2343** | **20** | **3** |
| **-** | **D-B** | **11:9** | **181** | **0** | **0** |
| **-** | **C-A** | **4:4** | **56** | **0** | **0** |

The residues forming inter-protomer hydrogen bonds are spread across multiple secondary structure elements of ArnC, including β3, β3-α3 loop, α7-α8 loop, α8, IH2, TM1, TM2, C-linker, and β8. Similar bonding patterns are observed for both interfaces, which includes interactions between β3 strand (S71 and L73) and β8 (Q311, Q312, I314), and between β3-α3 loop (N75, R76) and C-linker (F309), as well as IH1 (T151/T152). Further, interactions between R284 residues of both TM2 helices, salt bridges between D223 of IH2 and R234 of TM1, and hydrogen bonding between R180 located on the α7-α8 loop of the D/B protomer and D301 of TM2, as well as R305 and R307 of the C-linker are also shared in both interfaces.

While both type of interfaces have comparable biophysical properties and share the general pattern of interactions, specific differences result in C2 symmetry for the tetrameric assembly. Exclusively in interface type I (in both DA and BC dimers), the β3-α3 loop (75 to 78) of the D/B protomer interacts with the α8 helix (183 to 192), where R76 [NH1] coordinates hydrogen bonding with the backbone atoms in N189, I190, and A192, N75 interacts with F191, and Y78 forms a hydrogen bond with F191. The α8 helix is followed by the β7 strand, which leads to IH2 (α9), and its orientation could be modulated by IH2 conformational changes.

Additionally, there are also differences in interactions between IH2, TM1, and TM2 in the interfaces are noted. In interface type I, R234 on TM1 of the A/C protomer forms a salt bridge with D223 on IH2 of the D/B protomer. In contrast, in interface type II, R234 on TM1 of the B/D protomer forms three salt bridges with D223 and a hydrogen bond with N219 located on IH2 of the A/C protomer Further, there is an additional hydrogen bond between S237 of TM1 and Y222 of IH2. Conversely, interface type I features stronger interactions between IH2 and TM2, including an exclusive salt bridge between D223 of IH2 and R297 of TM2. Collectively, these interactions spaced along each of the protomers stabilize the ArnC tetramer and fix the relative positioning of the GT domain, the IH1/IH2 region, and the TM region.

### 4.3 Structure of UDP bound ArnC

ArnC catalyzes the transfer of UDP-L-Ara4N from the cytosol to produce UndP-Ara4FN in the inner membrane. To gain insights into the mechanism of reaction catalyzed by ArnC, we performed cryo-EM SPR of ArnC incubated with 1 mM Mn^2+^ and 1 mM UDP. The analysis of obtained the structure refined to 3.8 Å revealed a ligand-bound state of ArnC with A-loop, spanning 201 to 213, in the closed conformation (Figure 3A). The UDP binding pocket involves 14 amino acid residues from the ArnC IH1/2 and A-loop and suggests the location of the sugar-binding site in this protein (Figure 3B). UDP binding resulted in conformational changes, reflected by the RMSD of 2.46 Å and TM-score of 0.88, between the bound and unbound structures of ArnC. Apart from stabilization of the A-loop, which is postulated to be involved in ArnC catalytic activity, these changes also involved the position of IH1 and IH2 (Figure 3A). Specifically, in the UDP bound structure, the IH2 shifted towards the UDP binding pocket. Overall, UDP binding symmetrizes the tetrameric arrangement as more similarities are observed in interfaces between ArnC protomers.

**Figure 3:**
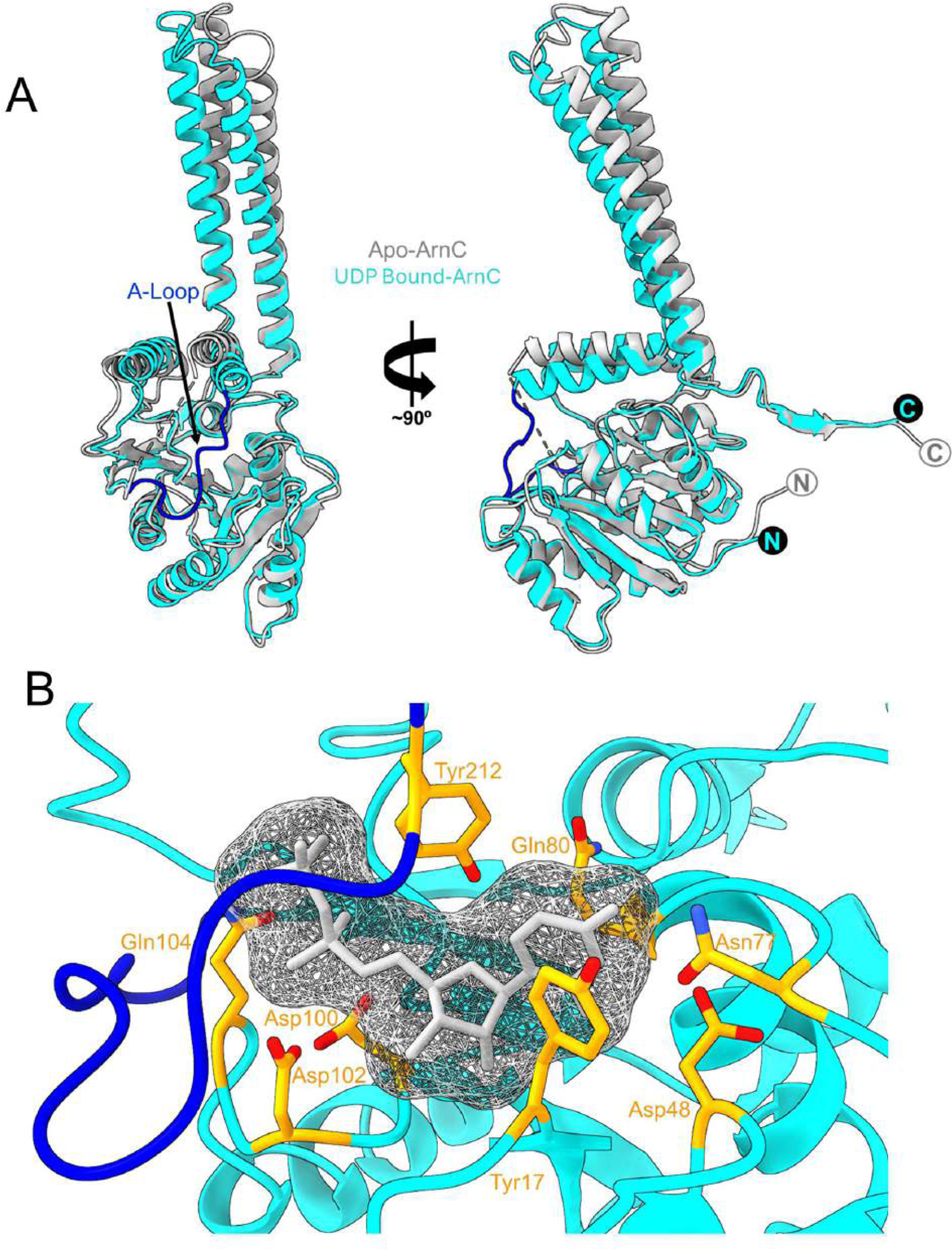
Structure of ArnC tetramer (A) Structural alignment of apo and UDP bound ArnC. (B) UDP binding pocket formed by IH1, IH2, and A-loop.

The GT-2 family of enzymes features a conserved DxD motif that binds divalent cations to coordinate the diphosphate group of the sugar donor (30). In ArnC, this motif corresponds to residues ^100^DADLQ^104^. However, the low resolution of the reconstruction and partial disorder between G207 to K211 prevented the modeling of the divalent cation’s position in the ArnC-UDP complex structure.

### 4.4 Comparison to structural homologs

To gain insight into catalytically important features of ArnC, we compared the obtained structure of ArnC to the two structurally characterized membrane-bound GT-2 enzymes - DPMS (Dolichol Phosphate Mannose Synthase) from *Pyrococcus furiosus* (PDB:5MLZ) and GtrB (PDB: 5EKP) from *Synechocystis* (28, 29). The structure of the ArnC protomer is most similar to that of GtrB (Figure 4A). These protomers share a similar topology and can be superimposed with a RMSD of 3.5 Å across 307 Cα atoms. However, the lack of electron densities in the GtrB crystal structure left the conformation of its periplasmic loop undefined; thus, restricting the interpretation of the topology of TM2 and the role of the C-terminal β-strands in intermolecular interactions (28). According to our structural analysis, the C-terminal β8 strand is involved in stabilizing the ArnC tetramer. The observed general similarities between ArnC and GtrB proteins suggest a comparable oligomeric arrangement for the latter with respect to the C-terminal structural elements.

**Figure 4:**
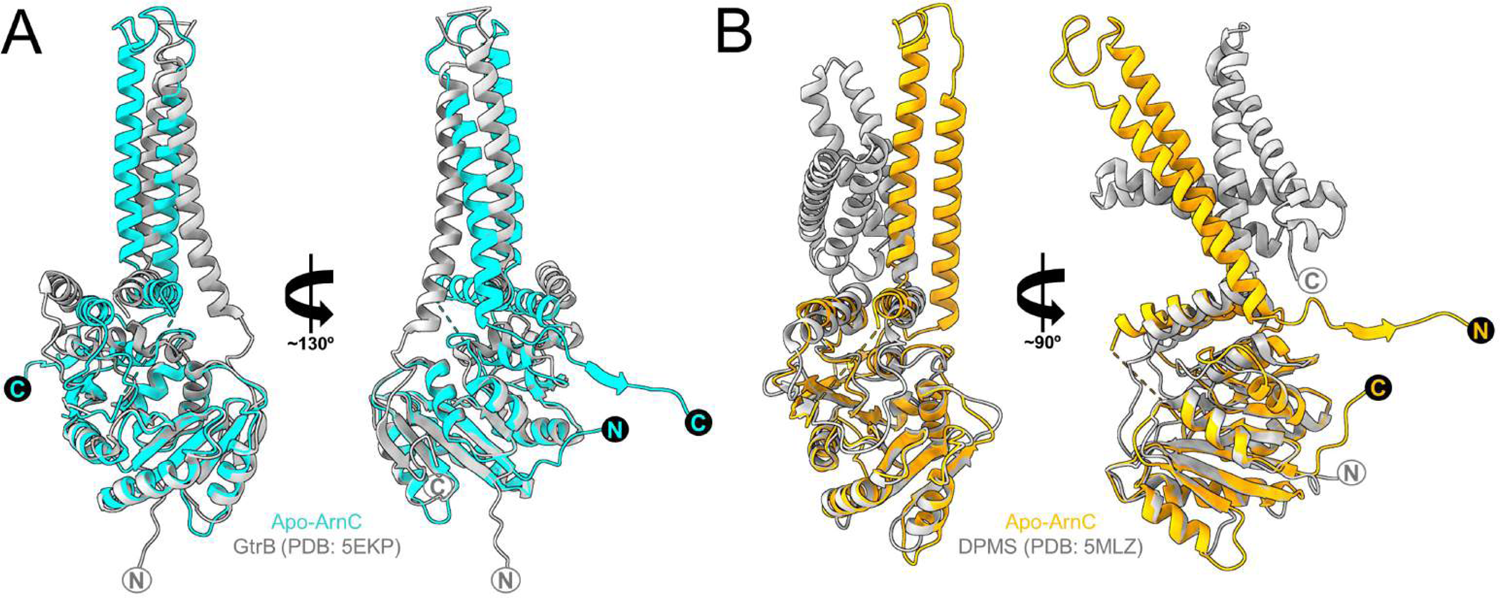
Structural comparison with GT-2 homologs. (A) Structural alignment of ArnC and GtrB protomers (B) Structural alignment of ArnC and DPMS protomers.

Comparison between DPMS and ArnC structures highlighted significant differences in the architecture of the transmembrane region in these proteins despite 28% of primary sequence identity between these two proteins (Figure 4B). While ArnC features two extended TM helices, DPMS has four TM helices that anchor this protein to the endoplasmic reticulum (ER) membrane. Additionally, the DPMS is expected to function as a monomer or dimer, unlike the tetrameric ArnC.

The cytoplasmic GT-2 domains in DPMS and ArnC share some common molecular features despite differences in substrate specificities. The residues D91 and Q93 forming the DxD motif in the DPMS catalytic pocket correspond to D102 and Q104 in ArnC, further supporting suggested role on the latter pair in sugar substrate coordination (Figure 5). Based on structural similarity and primary sequence analysis, ArnC residue R205 on the A-loop corresponds to a residue in DPMS involved in the coordination of the diphosphate of the sugar carrier (Figure 5A). Along the same lines, ArnC residues S141, R128, and R137 located on IH1 can be involved in the coordination of the phosphate head group of the lipid substrate, while residues A140, I144 and I148 on IH1, and F214 and M215 on IH2 can be involved in interactions with the first two isoprene units (Figure 5B). Notably, the conformational change in orientation of IH2 upon UDP binding observed in ArnC was also reported in the case of DPMS. Accordingly, we hypothesize that ArnC and DPMS have similar catalytic mechanisms where the A-loop in these enzymes gates the access to the catalytic site, which is modulated by changes in conformation of IH2.

**Figure 5:**
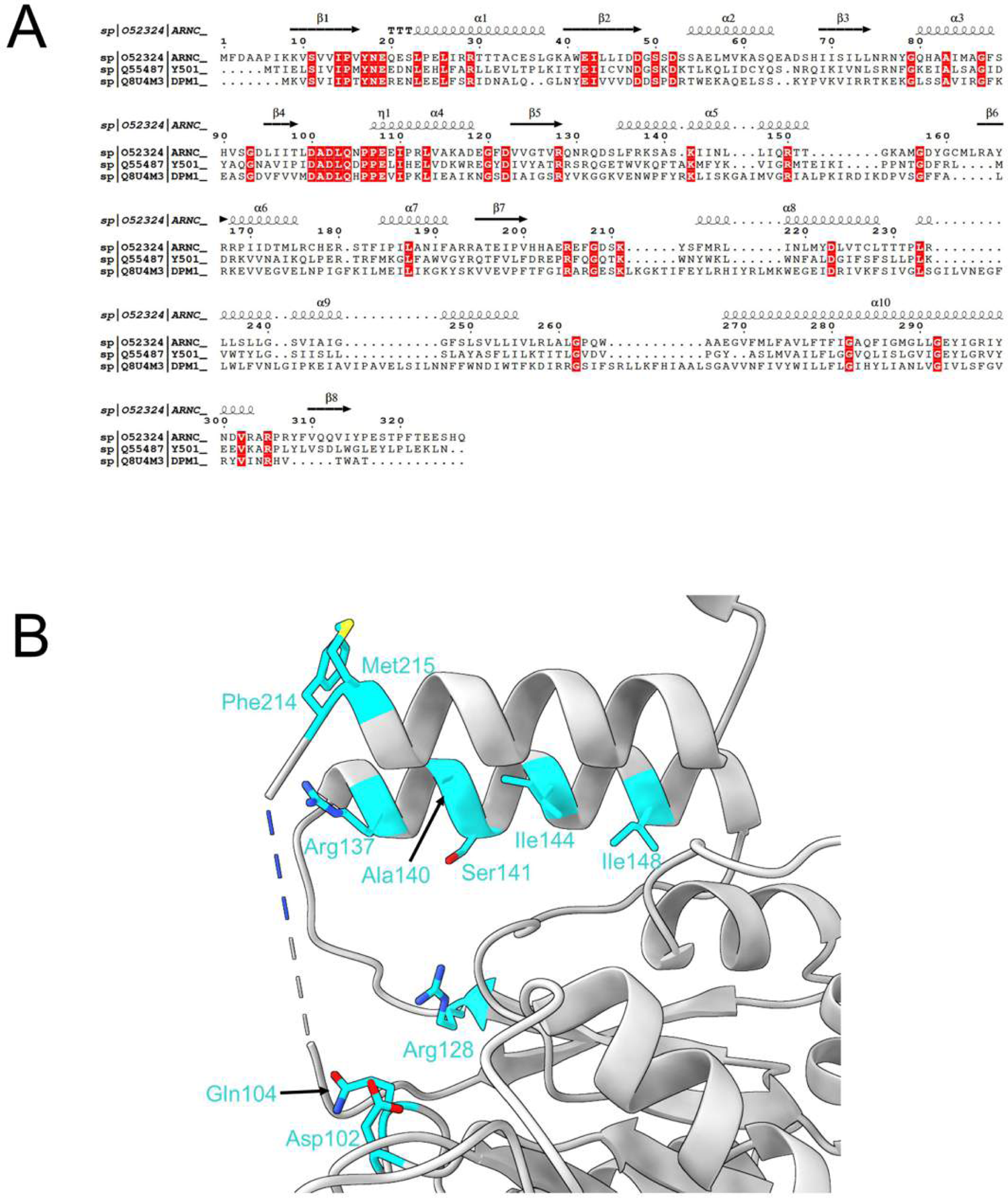
Putative catalytic residues. (A) Primary sequence alignment of ArnC, GtrB, and DPMS (B) Putative catalytic residues in ArnC involved in binding sugar donor and phospholipid acceptor.

## 5. Conclusion

This study presents the first experimentally derived model of integral membrane protein ArnC from lipid-A modification pathway involved in developing polymyxin resistance in Gram-negative pathogens. The cryo-EM structure of ArnC in apo form determined at 2.75 Å resolution the three main structural elements of this protein’s protomer and defines the mechanism of its tetrameric arrangement. The cryo-EM reconstruction of ArnC in UDP bound form provided insight on the location of the substrate binding site and defines the role of previously unresolved A-loop as a catalytically important structural element. Based on our comparative analysis with homologues, GtrB and DPMS, we are able to suggest other residues in ArnC that could be involved in its catalytic activity, providing the roadmap for future functional characterization.

## Author Contributions

DHP: Conceptualization, formal analysis, investigation, methodology, validation, visualization, writing – original draft preparation, writing – review and editing. EK: Conceptualization, investigation, methodology, writing – review and editing. YG: Conceptualization, formal analysis, investigation, methodology, validation, visualization, writing – original draft preparation, writing – review and editing. CS: Writing – original draft preparation. DTP: Visualization, writing – review and editing. TE: Investigation. DB: Conceptualization, formal analysis, investigation, methodology, validation, funding acquisition, supervision, writing – original draft preparation, writing – review and editing. AS: Conceptualization, formal analysis, investigation, methodology, validation, funding acquisition, supervision, visualization, writing – original draft preparation, writing – review and editing.

## Acknowledgments

We thank the members of Savchenko lab for insightful discussions. This study was funded in whole or in part by the U.S. Federal funds from the National Institute of Allergy and Infectious Diseases (Contract 75N93022C00035) and structures were solved as part of the Center for Structural Biology of Infectious Diseases (CSBID). We thank Dr. Jennifer Kohler and her lab members, Mary Burns, B.S., and Emanuela Capota, B.S., for providing the 10 mM stock solution of UDP. D.T.P was supported by the Doctoral Alberta Graduate Excellence Scholarship. We thank the Cryo-Electron Microscopy Facility (CEMF) at UT Southwestern Medical Center which has been supported by grants RP220582 from the Cancer Prevention and Research Institute of Texas (CPRIT) for maintaining microscopes. Some data presented in this report were acquired with a mass photometer that was supported by award S10OD030312-01 from the National Institutes of Health.

---

**Figure 1S.**
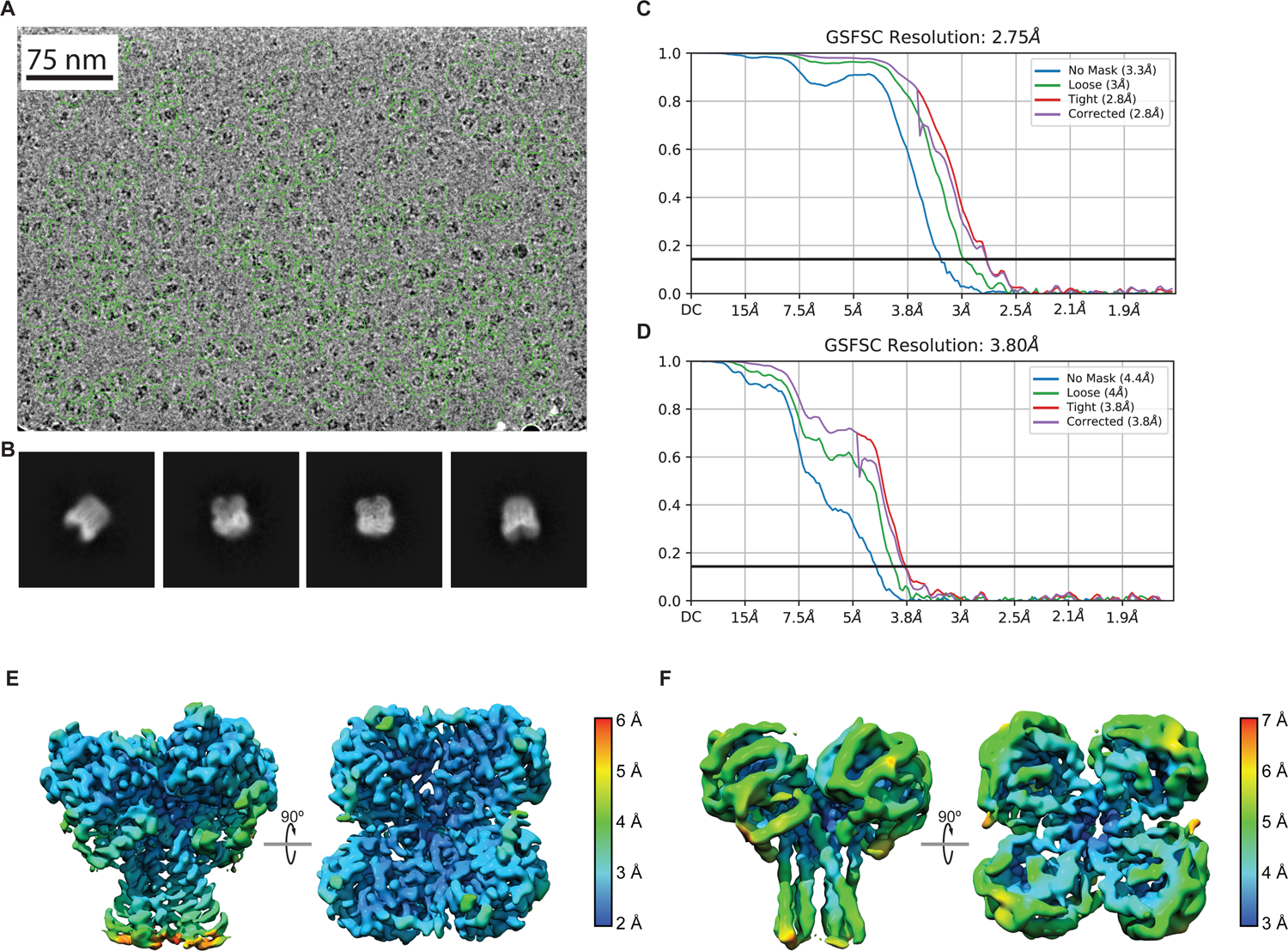
Selected cryo-EM workflow results A) Motion corrected micrograph for ArnC apo. Particles are shown in green circles. B) Selected 2D classes for ArnC apo. Micrographs and particles of ArnC+UDP+Mn^2+^ are similar to those of ArnC apo and thus are not shown. C) FSC curve of the final ArnC apo reconstruction. D) FSC curve of the final ArnC+UDP+Mn^2+^ reconstruction. E) Local resolution of ArnC apo reconstruction. F) Local resolution of ArnC+UDP+Mn^2+^ reconstruction.

